# Deconvolution of DNA methylation signatures identifies common differentially methylated gene regions on 1p36 across breast cancer subtypes

**DOI:** 10.1101/115873

**Authors:** Alexander J. Titus, Gregory P. Way, Kevin C. Johnson, Brock C. Christensen

## Abstract

Breast cancer is a complex disease and studying DNA methylation (DNAm) in tumors is complicated by disease heterogeneity. We compared DNAm in breast tumors with normal-adjacent breast samples from The Cancer Genome Atlas (TCGA). We constructed models stratified by tumor stage and PAM50 molecular subtype and performed cell-type reference-free deconvolution on each model. We identified nineteen differentially methylated gene regions (DMGRs) in early stage tumors across eleven genes (*AGRN, C1orf170, FAM41C, FLJ39609, HES4, ISG15, KLHL17, NOC2L, PLEKHN1, SAMD11, WASH5P*). These regions were consistently differentially methylated in every subtype and all implicated genes are localized on chromosome 1p36.3. We also validated seventeen DMGRs in an independent data set. Identification and validation of shared DNAm alterations across tumor subtypes in early stage tumors advances our understanding of common biology underlying breast carcinogenesis and may contribute to biomarker development. We also provide evidence on the importance and potential function of 1p36 in cancer.

## INTRODUCTION

Invasive breast cancer is a complex disease characterized by diverse etiologic factors^1^. Key genetic and epigenetic alterations are recognized to drive tumorigenesis and serve as gate-keeping events for disease progression^2^. Early DNA methylation (DNAm) events have been shown to contribute to breast cancer development^3^. Importantly, DNAm alterations have been implicated in the transition from normal tissue to neoplasia^4,5^ and from neoplasia to metastasis^6^. Furthermore, patterns of DNAm are known to differ across molecular subtypes of breast cancer^7^ - Luminal A (LumA), Luminal B (LumB), Her2-enriched and Basal-like - identified based on the prediction analysis of microarray 50 (PAM50) classification^8^. However, while DNAm differences across breast cancer subtypes have been explored, similarities across subtypes are less clear^9^. Such similarities found in early stage tumors can inform shared biology underpinning breast carcinogenesis and – as similarities would be agnostic to subtype – potentially contribute to biomarkers for early detection.

Studying DNAm in bulk tumors is complicated by disease heterogeneity. Heterogeneity is driven by many aspects of cancer biology including variable cell-type proportions found in the substrate used for molecular profiling^10^. Different proportions of stromal, tumor, and infiltrating immune cells may confound molecular profile classification when comparing samples^11^ because cell types have distinct DNAm patterns^12–14^. The potential for cell–type confounding prompted the development of statistical methods to adjust for variation in cell-type proportions in blood^15^ and solid tissue^16^. One such method, *RefFreeEWAS*, is a reference-free deconvolution method and does not require a reference population of cells with known methylation patterns and is agnostic to genomic location when performing deconvolution^17^. Instead, the unsupervised method infers underlying cell-specific methylation profiles through constrained non-negative matrix factorization (NMF) to separate cell-specific methylation differences from actual aberrant methylation profiles observed in disease states. This method has previously been shown to effectively determine the cell of origin in breast tumor phenotypes^18^.

We applied *RefFreeEWAS* to The Cancer Genome Atlas (TCGA) breast cancer DNAm data and estimated cell proportions across the set. We compared tumor DNAm with adjacent normal tissue stratified by tumor subtype^9^ and identified common early methylation alterations across molecular subtypes that are independent of cell type composition. We identified a specific chromosomal location, 1p36.3, that harbors all 19 of the differentially methylated regions that are in common to early stage breast cancer subtypes. 1p36 is a well-studied and important region in many different cancer types, but there remain questions about how it may impact carcinogenesis and disease progression^19^. Our study provides evidence that methylation in this region may provide important clues about early events in breast cancer. We also performed *RefFreeEWAS* on an independent validation set (GSE61805) and confirmed these results^20^.

## RESULTS

### DNA methylation deconvolution

Subject age and tumor characteristic data stratified by PAM50 subtype and stage is provided in Table 1 for the 523 TCGA tumors analyzed. TCGA breast tumor sample purity, estimated by pathologists from histological slides, was consistent across PAM50 subtypes and stages indicating that observed methylation differences are not predominantly a result of large differences in tumor purity (Supplementary Fig. S1). To correct for cell-proportion differences across tumor samples, we estimated the number of cellular methylation profiles contributing to the mixture differences by applying NMF to the matrix of beta values, which resulted in model specific dimensionality estimates indicating diverse cellular methylation profiles (Supplementary Table S1). The reference-free deconvolution altered the number and extent of significant differentially methylated CpGs across all models that compared breast tumor methylation with adjacent normal samples (Supplementary Fig. S2).

**Table 1.** Sample information stratified by PAM50 subtype.

|  | Basal-like | Her2 | Luminal A | Luminal B | Total with Assignment |
| --- | --- | --- | --- | --- | --- |
| TCGA tumors | 86 | 31 | 279 | 127 | 523 |
| Age, mean (SD) | 56.8 (12.8) | 60 (12.8) | 58 (13.5) | 57.1 (12.6) | 57.8 (13.1) |
| Stage*, n (%) | -- | -- | -- | -- | -- |
| Early (I/II) | 70 (81%) | 20 (65%) | 207 (74%) | 84 (66%) | 381 (73%) |
| Late (III/IV) | 14 (16%) | 10 (32%) | 69 (25%) | 42 (33%) | 135 (26%) |
| Missing | 2 (2%) | 1 (3%) | 3 (1%) | 1 (1%) | 7 (1%) |

\*AJCC characterized stage, provided by TCGA

### Subtype specific methylation patterns

In early stage tumors, we identified a set of nineteen DMGRs shared among Luminal A, Luminal B, Her2, and Basal-like subtypes (DMGRs *Q* < 0.01, Figure 1A). In the late stage tumors, we identified 31,931 DMGRs in common across subtypes (Figure 1B). Subtype specific methylation patterns in early stage tumors were most divergent for Basal-like tumors versus other types, while in late stage tumors methylation alterations in Luminal B tumors were most divergent (Supplementary Table S2). To test if collapsing by genomic region had an appreciable effect on detecting DMGRs, we compared DMGR results to results derived from regions defined by CpG island status (i.e. CpG island, Shore, Shelf, Open Sea). Using CpG island context designations indicated similar results (Supplementary Fig S3), though a lower number of common DMGRs were observed. Therefore, downstream analyses used DMGRs identified based on probe position in relation to TSS.

**Figure 1.**
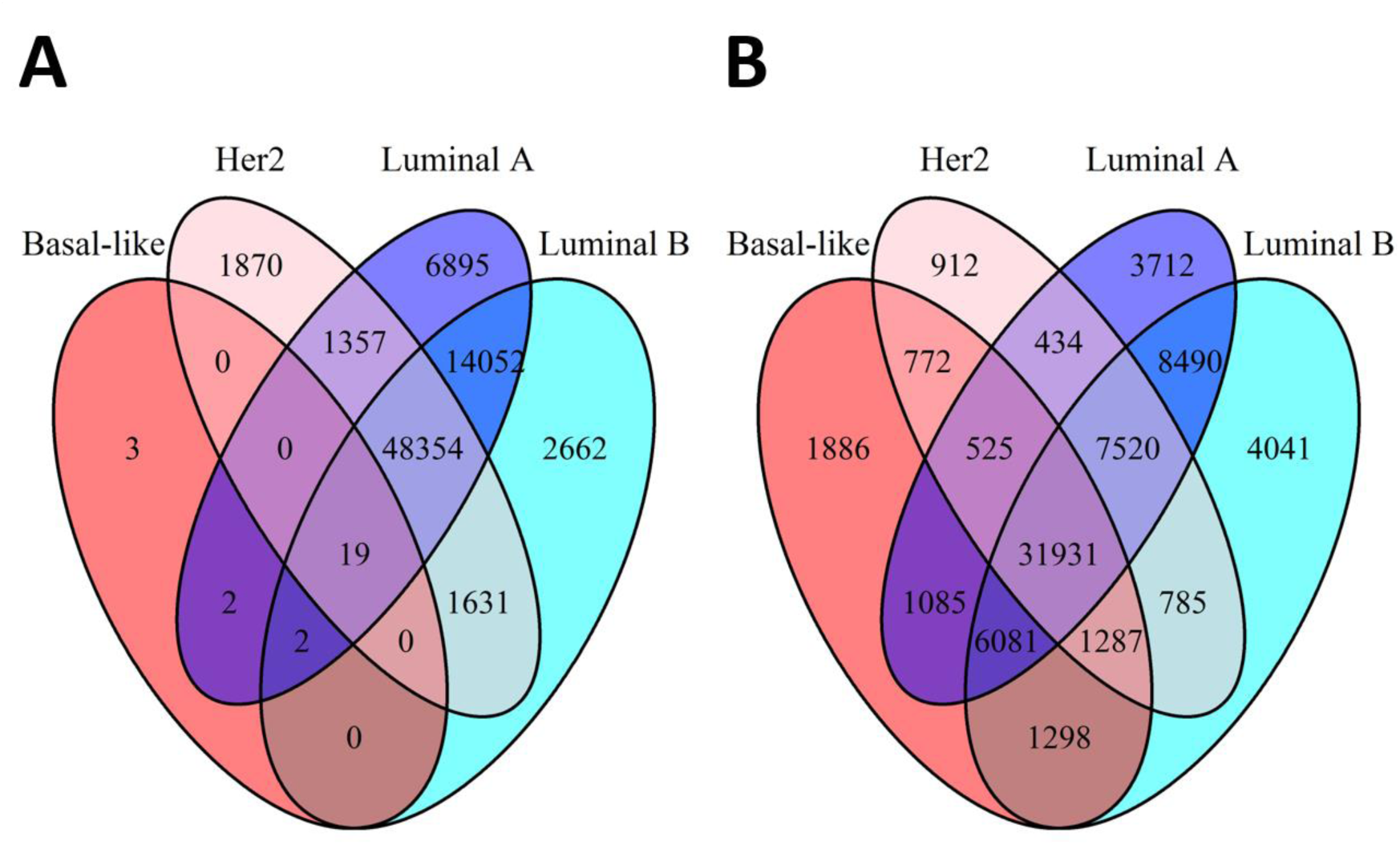
Numbers of overlapping differentially methylated gene regions in (A) early stage tumors (n = 76,847) and (B) late stage tumors (n = 70,759) stratified by Basal-like, Her2, Luminal A, and Luminal B PAM50 subtypes with a Q-value cutoff of 0.01.

We identified nineteen DMGRs with common methylation alterations among tumor subtypes in comparison with normal tissues that were annotated to eleven genes: *AGRN, C1orf170, FAM41C, FLJ39609, HES4, ISG15, KLHL17, NOC2L, PLEKNH1, SAMD11*, and *WASH5P* (Supplementary Table S3).

Dependent upon tumor subtype, some gene regions had a different directional change in tumor methylation compared to normal tissue (e.g. *C1orf170, HES4, and ISG15*). Additionally, of the eleven genes identified, we observed differential methylation in different regions including gene body, promoter (TSS1500, and TSS200), and 3’UTR (Table 2 and Supplementary Table S3). All nineteen DMGRs also had differential methylation in at least one late stage tumor subtype, and thirteen of the nineteen DMGRs were significantly differentially methylated across all tumor subtypes in late stage tumors (Table 2 and Supplementary Table S4). A heatmap of the unadjusted beta values for individual CpGs from the nineteen DMGRs demonstrated grouping of most of the Basal-like tumors separate from a group of mixed Luminal and Her2 tumors (Figure 2).

**Figure 2.**
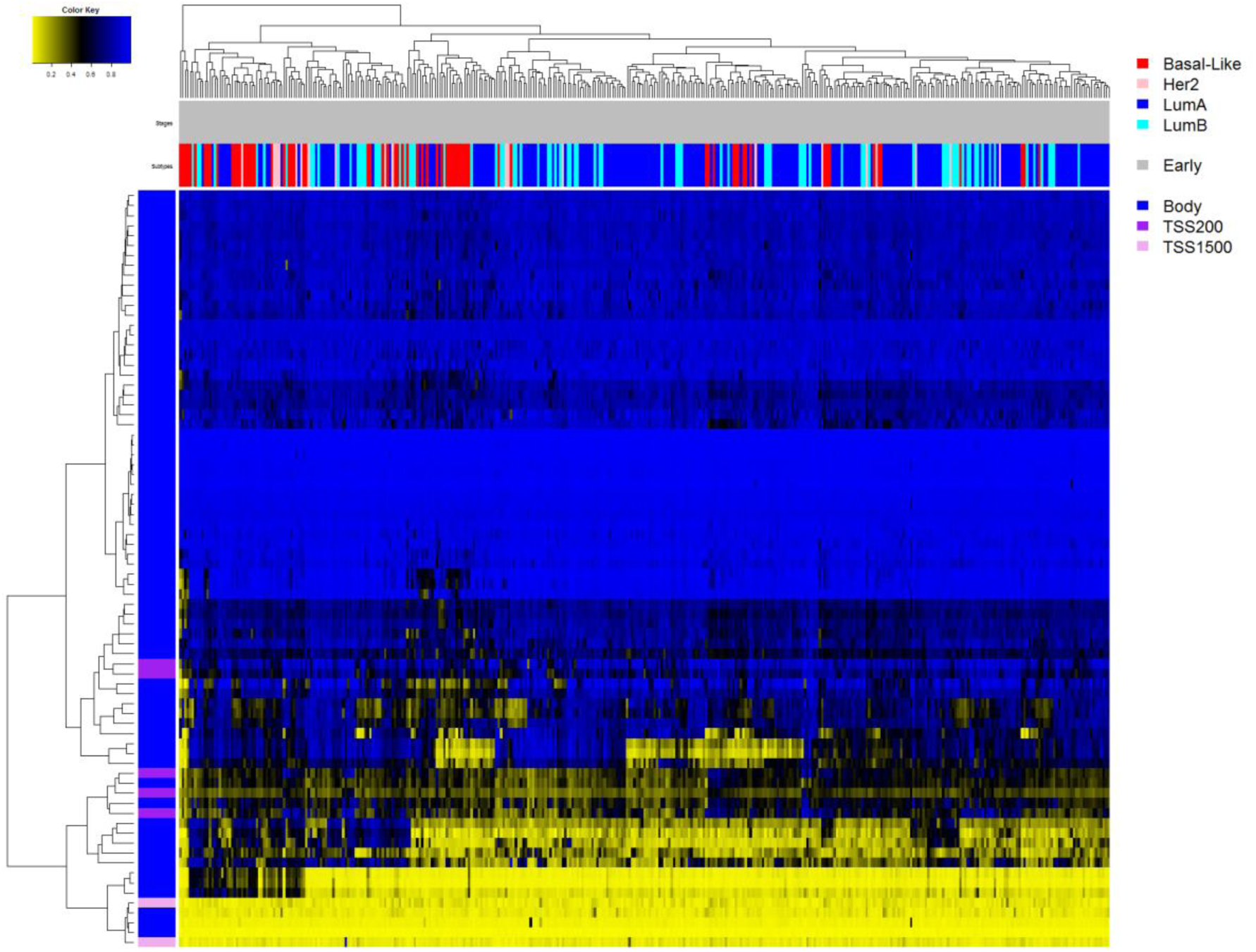
Raw beta value (unadjusted for cellular composition) heatmap of the significantly differentially methylated CpG sites mapping to the common early stage differentially methylated gene regions. The genomic context is given in the vertical color bar and the PAM50 subtype and tumor information (stage and subtype) are given in the horizontal bars. Yellow indicates low methylation and blue indicates high methylation beta values.

**Table 2.** Ninteen differentially methylated gene regions in common to early stage tumors.

| *DMGR* | *Alternate*<br>*Gene Name* | *Basal*<br>*Med Q* | *Her2*<br>*Med Q* | *Lum A*<br>*Med Q* | *Lum B*<br>*Med Q* | *\*Any late*<br>*stage* | *\*All late*<br>*stage* | *Present in*<br>*validation* | *Validation*<br>*Median Q* |
| --- | --- | --- | --- | --- | --- | --- | --- | --- | --- |
| AGRN Body | AGNR | 2.44E-06 | 1.68E-04 | 1.82E-07 | 1.29E-06 | Y | -- | Y | 7.80E-21 |
| C1orf170 Body | PERM1 | 4.03E-11 | 1.68E-05 | 5.46E-09 | 9.69E-04 | Y | Y | Y | 1.31E-08 |
| C1orf170 TSS1500 | PERM1 | 5.40E-04 | 6.52E-03 | 7.82E-06 | 6.82E-05 | Y | -- | Y | 9.23E-03 |
| FAM41C Body | FAM41C | 4.13E-03 | 4.20E-08 | 1.18E-20 | 3.43E-03 | Y | Y | Y | 8.25E-10 |
| FAM41C TSS1500 | FAM41C | 3.27E-04 | 1.11E-04 | 8.38E-05 | 1.04E-34 | Y | Y | Y | 1.75E-24 |
| FLJ39609 TSS200 | LOC100130417 | 1.30E-04 | 6.02E-05 | 2.92E-06 | 3.67E-04 | Y | Y | Y | 5.24E-06 |
| HES4 TSS1500 | HES4 | 3.06E-03 | 5.15E-04 | 7.84E-05 | 2.20E-04 | Y | -- | Y | 5.06E-04 |
| ISG15 Body | ISG15 | 3.14E-07 | 2.40E-04 | 1.18E-05 | 3.58E-04 | Y | Y | -- | 1.03E-01 |
| KLHL17 3'UTR | KLHL17 | 3.14E-05 | 5.51E-07 | 3.83E-16 | 2.27E-03 | Y | Y | Y | 3.99E-08 |
| KLHL17 Body | KLHL17 | 5.90E-06 | 1.10E-04 | 7.85E-04 | 7.24E-05 | Y | -- | Y | 1.60E-06 |
| NOC2L Body | NOC2L | 3.15E-04 | 6.15E-04 | 6.56E-05 | 2.40E-06 | Y | Y | Y | 4.90E-11 |
| PLEKHN1 3'UTR | PLEKHN1 | 5.17E-16 | 4.73E-06 | 3.10E-07 | 7.74E-06 | Y | -- | Y | 9.83E-09 |
| PLEKHN1 Body | PLEKHN1 | 8.94E-10 | 2.71E-09 | 7.58E-29 | 1.73E-30 | Y | Y | Y | 5.87E-18 |
| PLEKHN1 TSS1500 | PLEKHN1 | 3.14E-05 | 5.51E-07 | 2.59E-06 | 3.63E-07 | Y | Y | Y | 3.99E-08 |
| PLEKHN1 TSS200 | PLEKHN1 | 1.56E-18 | 5.77E-10 | 1.42E-03 | 1.22E-03 | Y | Y | Y | 2.93E-10 |
| SAMD11 5'UTR | SAMD11 | 3.58E-03 | 7.23E-12 | 1.01E-09 | 2.21E-08 | Y | Y | Y | 4.59E-11 |
| SAMD11 Body | SAMD11 | 7.13E-08 | 2.47E-08 | 8.49E-06 | 2.04E-04 | Y | Y | Y | 3.26E-23 |
| SAMD11 TSS1500 | SAMD11 | 2.38E-03 | 6.14E-04 | 8.56E-04 | 1.02E-03 | Y | Y | Y | 2.02E-05 |
| WASH5P Body | WASH7P | 2.93E-03 | 9.84E-03 | 1.64E-03 | 1.25E-05 | Y | -- | -- | 7.01E-02 |

\*Reference to any or all breast cancer subtypes in late stage tumors

### DMGRs cluster on chromosome 1p36 and on gene bodies

Of the nineteen DMGRs identified, all of them are in eleven genes located on the *p36.3* cytoband of chromosome 1 (Supplementary Figure S4). Chromosome 1p36.3 is the start section of chromosome 1 and of the eleven genes identified, one *(WASH5P)* is located near the very start of the chromosome (chr1:14,362 - 29,370) and the other ten genes are located end-to-end between chr1:868,071 - 1,056,116 (Supplementary Figure S4).

Most of the DMGRs tracked to gene body regions: *AGRN, C1orf170, FAM41C, ISG15, KLHL17, NOC2L, PLEKHN1, SAMD11*, and *WASH5P* all had gene body methylation differences. Gene body regions were enriched among early stage tumor DMGRs compared to all other regions (Fisher’s Exact Test OR = 4.15, 95% CI = 1.04 – 23.83, *P* = 0.04). All differentially methylated CpG probe IDs are given in Supplementary Table S5. DAVID pathway analysis applied to the top 400 most aberrantly methylated genes in common to the four PAM50 subtypes identified the GO term for the regulation of hormone levels to be significantly enriched (GO:0010817, *FDR* = 0.035, Supplementary Table S6).

### Breast cancer copy number alterations in 1p36

Among these 523 tumors, the prevalence of 1p36.3 copy number alteration was only 1.2% (n=6), all were amplifications that affected ten of the eleven genes most distal to the chromosome end. Among the six tumors with 1p36.3 amplification three were Basal-like, two were Her2-enriched, and one was Luminal A. Exclusive of tumors with copy number alterations, there was one tumor (Her2-enriched), with a truncating mutation in *KLHL17*, and one tumor with a missense mutation in *PLEKHN1* (Basal-like).

### DMGRs impact gene expression

We identified CpG sites with significant correlation of methylation with gene expression for five genes (*AGRN, PLEKHN1, KLHL17, SAMD11*, and *FAM41C*), associated with eight DMGRs (Supplementary Table S7 and Supplementary Figures S6-9).

### Validating DMGR hits in an independent dataset

We validated our findings in an independent 450K methylation data set from 186 tumors and 46 normal tissues described in Fleischer *et al.* (GSE60185). Seventeen of nineteen DMGRs were significantly differentially methylated between tumor and normal tissues in the replication set (all DMGRs at *Q* < 0.01; Table 2), and CpGs in these DMGRs had similar patterns of beta value distributions (Supplemental Figure S10). The remaining two gene regions were also highly ranked in the *q* value distribution (*WASH5P* body: *Q* = 0.07; *ISG15* Body: *Q* = 0.10).

### Reproducibility

All TCGA and validation data is publicly available. We also provide software under an open source license for analysis reproducibility and to build upon our work^21^.

## DISCUSSION

We were interested in identifying common biology underlying breast cancer independent of molecular subtype and cell-type proportion. After applying a reference-free deconvolution algorithm, we observed that early stage tumors harbor differentially methylated gene regions localized entirely to a small region on 1p36.3 shared across four major subtypes. Although DNA methylation alterations are widespread in early stage tumors and prior work has demonstrated alterations that differ among breast tumor subtypes^9,22^ we observed only 19 DMGRs that overlapped molecular subtypes. All DMGRs tracking to the same region on 1p36.3 suggests that altered regulation of this region contributes to breast carcinogenesis irrespective of disease subtype.

Previously, alterations on chromosome 1 have been observed in breast cancer cell lines and tumors^23^. Additionally, copy number deletions in this region have been shown to be an important precursor in DCIS tumors ^24^ and in follicular lymphomas ^25^. However, the most prevalent copy number alterations on chromosome 1 are gains on the *q* arm and losses on the *p* arm that do not typically fully encompass our implicated genes on 1p36.3^23,26,27^. The region is also well-studied and significantly altered in neuroblastoma - the most common solid tissue tumor of childhood^28–31^. The biological underpinnings of this region remain elusive^19,32^ but a systematic understanding of how these specific DMGRs may impact early cancer development may be important for other cancer types and not just breast cancer.

Of the nineteen DMGRs identified, eighteen of them replicated in either one or both late stage and independent validation sets. The one DMGR that did not replicate was the *WASH5P body.* This region is located more than 830,000 base pairs (bp) away from the much tighter region spanned by the remaining eighteen DMGRs (~188,000 bp), suggesting a loose association between *WASH5P* and the other ten genes.

There is also additional evidence implicating the potential importance of the identified genes assigned to the differentially methylated regions. For example, in a study of mutational profiles in metastatic breast cancers, *AGRN* was more frequently mutated in metastatic cancers compared with early breast cancers^33^. Similarly, expression of the *HES4* Notch gene is known to be significantly correlated with the presence of activating mutations in multiple breast cancer cell lines, and is associated with poor patient outcomes^34^. In addition, *ISG15* has been implicated as a key player in breast carcinogenesis^35^, though there is conflicting evidence^36^. However, the conflicting evidence to date may be related to our observation of *ISG15* hypomethylation in Basal-Like, Her2, and LumB tumors, and hypermethylation in LumA tumors (Supplementary Table S3). Opposing methylation states among tumor subtypes relative to normal tissue may contribute to subtype-specific roles of *ISG15* dysregulation in breast carcinogenesis. Additionally, the *NOC2L* gene has been identified as a member of a group of prognostic genes derived from an integrated microarray of breast cancer studies^37^. We also identified three DMGRs – TSS1500, Body, & 5’UTR - in the *SAMD11* gene, which has significantly reduced expression in breast cancer cells compared to normal tissues^38^, consistent with our findings of *SAMD11* hypermethylation across all four breast cancer subtypes. As DNAm changes were observed consistently and robustly across subtypes, it is likely that several of the other identified genes are cancer initiation factors that require additional study.

Importantly, we validated the identified DMGRs in an independent set of invasive breast tumors and normal tissues. Our validation is strengthened by the lack of molecular subtype assignments in the validation set. The validation of DMGRs in a setting agnostic to intrinsic subtype indicates that differential magnitude or direction of methylation alterations that may be present in different subtypes did not limit our ability to identify significant alterations. A limitation of the validation set is a lack of gene expression data to further investigate relationships between expression and methylation for each gene region. Nevertheless, additional targeted studies on this set of validated genes and gene regions can enhance the understanding of methylation alterations at these DMGRs in breast carcinogenesis.

Caution should be exercised in interpreting the results of the adjusted beta coefficients from the reference-free algorithm. It is unclear if specific disease states are a result of aberrant methylation profiles in specific cell types which then cause changes to cell mixtures, or if the disease state is a result of cell-type proportion differences. Additionally, the unsupervised clustering heatmaps plot unadjusted methylation beta values and do not account for cell type adjustment. Lastly, the DMGR analysis drops CpGs that do not track to gene regions, which may reduce detection of non-genic regions related with breast carcinogenesis.

We identified and validated DMGRs in early stage breast tumors across PAM50 subtypes that are located on chromosome 1p36.3. The observed differential methylation suggests that this region may contribute to the initiation or progression to invasive breast cancer. Additional work is needed to investigate the scope of necessary and sufficient alterations to 1p36.3 for transformation and to more clearly understand the implications of 1p36.3 methylation alterations to gene regulation. Further investigation of DNAm changes to 1p36.3 may identify opportunities for early identification of breast cancer or risk assessment. Lastly, the reference-free approach we used could be applied to methylation datasets from other tumor types to identify potential drivers of carcinogenesis common across histologic or intrinsic molecular subtypes.

## PATIENTS & METHODS

### Data Processing

We accessed breast invasive carcinoma Level 1 Illumina HumanMethylation450 (450K) DNAm data (n = 870) from the TCGA data access portal and downloaded all sample intensity data (IDAT) files. We processed the IDAT files with the R package *minfi* using the “Funnorm” normalization method on the full dataset ^39^. We filtered CpGs with a detection *P*-value > 1.0E-05 in more than 25% of samples, CpGs with high frequency SNP(s) in the probe, probes previously described to be potentially cross-hybridizing, and sex-specific probes ^40,41^. We filtered samples that did not have full covariate data (PAM50 subtype, pathologic stage^42,12^) and full demographic data (age and sex). All tumor adjacent normal samples were included regardless of missing data (n = 97, Table 1).

From an original set of 485,512 measured CpG sites on the Illumina 450K array, our filtering steps removed 2,932 probes exceeding the detection *P*-value limit, and 93,801 probes that were SNP-associated, cross-hybridizing, or sex-specific resulting in a final analytic set of 388,779 CpGs. From 870 TCGA breast tumors, we restricted to primary tumors with available PAM50 intrinsic subtype assignments of Basal-like (n = 86), Her2 (n = 31), Luminal A (n = 279), and Luminal B (n = 127), excluding Normal-like tumors due to limited sample size (n = 18). Lastly, we restricted the final total tumor set to only those with stage assignments resulting in a final analytic sample size of n = 523.

### Reference-free cell type adjustment modeling

We stratified samples by PAM50 subtype (Basal-like, Luminal A, Luminal B, Her2) and then by tumor stage dichotomizing as early (stage I and II tumors) and late (stage III and IV tumors)^42^, resulting in eight distinct models. To analyze DNAm differences between tumor and normal tissue and to adjust for effects of cellular heterogeneity across samples, we applied the reference-free deconvolution algorithm from the *RefFreeEWAS* R package to each model adjusting for age^16^. The method estimates the number of underlying tissue-specific cell methylation states contributing to methylation heterogeneity through a constrained variant of NMF^43^. Briefly, the method assumes the sample methylome is composed of a linear combination of the constituent methylomes. It decomposes the matrix of sample methylation values *(Y)* into two matrices (*Y = ΜΩ^Τ^*), where M is an *m x K* matrix of m CpG-specific methylations states for K cell types and Ω is a *nx K* matrix of subject-specific cell-types. *K* is selected via bootstrapping *K* = 2…10 and choosing the optimal *K* that minimizes the bootstrapped deviance. To correct for multiple comparisons, we converted all extracted *P*-values to *Q*-values using the R package *qvalue*^44^.

### Identifying differentially methylated gene regions

To understand the genomic regions with common DNAm alterations we grouped CpGs by gene and region relative to genomic location (transcription start site 1500 (TSS1500), TSS200, 3’ untranslated region (3’UTR), 5’UTR, 1^st^ exon, and gene body). We used this gene-region taxonomy to collapse differentially methylated CpGs, as defined by our *Q*-value cutoff, into specific differentially methylated gene regions (DMGRs). This extended the Illumina 450K CpG annotation file to allow for a given CpG to be associated with up to two genes depending on the proximity of the CpG site to neighboring genes (Figure 3).

**Figure 3.**
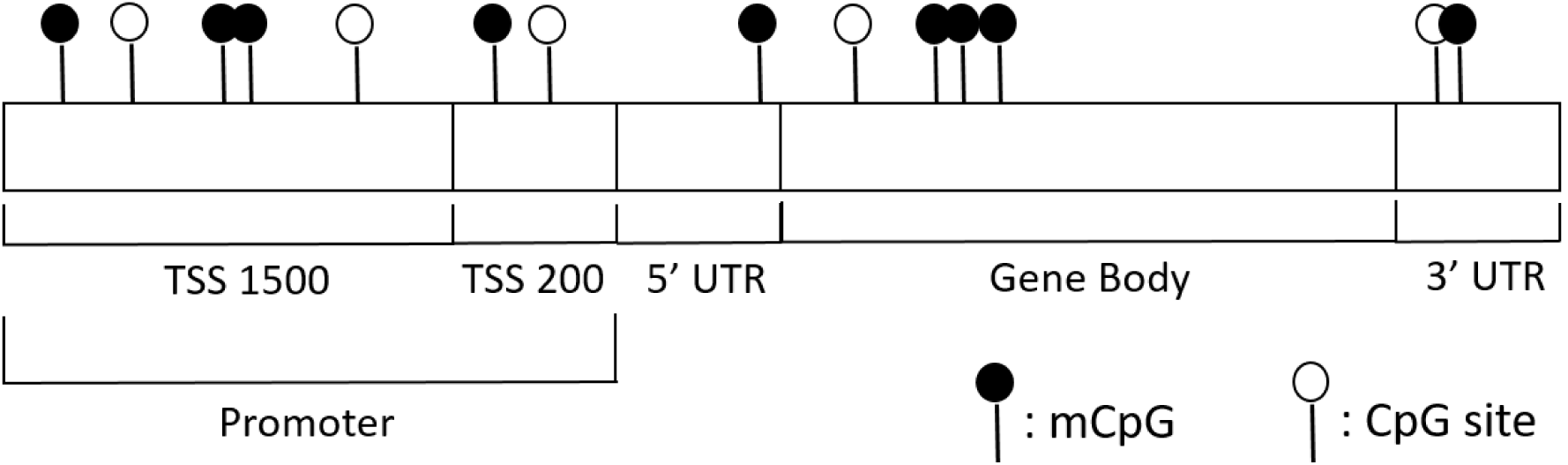
Diagram of_CpG sites relative to gene regions (Transcription start sites (TSS1500 & TSS200), Untranslated regions (5’UTR & 3’UTR), and the gene body). Dark circles indicate methylated sites and empty circles indicate unmethylated sites.

We defined a differentially methylated CpG as one with a Q-value < 0.01 following cell-type adjustment in a specific subtype model compared to normal tissue. To identify DMGR sets for each stage and subtype, we analyzed all eight models independently.

### Pathway Analysis

We performed a DAVID (the database for annotation, visualization and integrated discovery) analysis^45,46^ for the 400 genes with the lowest median CpG *Q*-values that are in common to all early stage tumors regardless of PAM50 subtype, and extracted enriched Gene Ontology (GO)^47^ and Kyoto Encyclopedia of Genes and Genomes (KEGG)^48^ terms. We selected the top 400 genes based on recommended gene list sizes^49^.

### Copy number, gene expression, and genomic location

We downloaded TCGA Breast Invasive Carcinoma CNV data^9^ and normalized RNAseq using cBioPortal^50^. For the DMGRs we identified, we analyzed the prevalence of copy number alterations and mutations in each gene across all samples, stratified by molecular subtype. Similarly, to determine whether these DMGRs affect gene expression of their target gene, we calculated Spearman correlations of DNAm beta values in significant CpGs (*Q* < 0.01) to matched sample Illumina HiSeq gene expression data. We used a Bonferroni correction to determine significant expression differences, resulting in an acceptance alpha value of 9.36E-5.

### Validation

To confirm the identified early stage DMGRs in common among intrinsic molecular subtypes we applied the analysis workflow to TCGA late stage tumors and an independent validation set (GSE60185)^20^. The validation set includes samples of ductal carcinoma *in situ* (DCIS), mixed, invasive, and normal histology collected from Akershus University Hospital and from the Norwegian Radium Hospital. We analyzed only the invasive samples compared to normal samples using the same bioinformatics pipeline of quality control CpG filtering steps and normalization procedures. However, we did not have complete age information or intrinsic subtype assignments for the validation set and the models are not adjusted for age or stratified by subtype. This resulted in a single model comparing 186 invasive tumors with 46 normal controls measured across 390,253 CpGs.

## ACKNOWLEDGEMENTS

Funding was provided by P20GM104416 and R01DE02277 (BCC), by the Quantitative Biomedical Sciences graduate program, and through a BD2K Fellowship to AJT (T32LM012204).

## SUPPLAMENTAL FIGURES

**Supplementary Figure S1.**
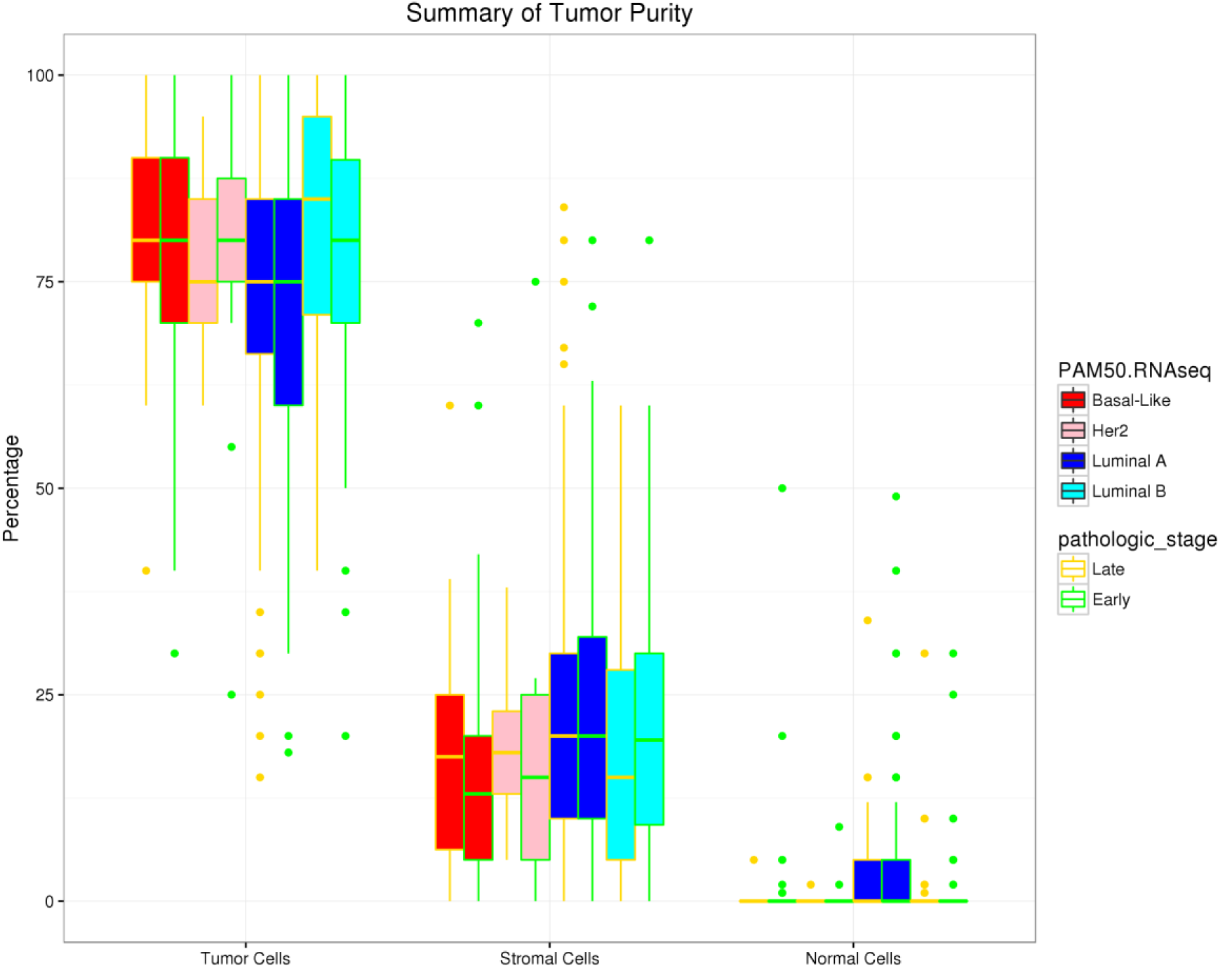
Box plots show the distribution of tumor purity across all subtypes for both early and late stages of the TCGA dataset. The measurements estimated by TCGA are based on histology slides and indicate the estimated distribution of the number of tumor cells, stromal cells, and normal cells in each sample. See the NCI CDE Browser for more details.

**Supplementary Figure S2.**
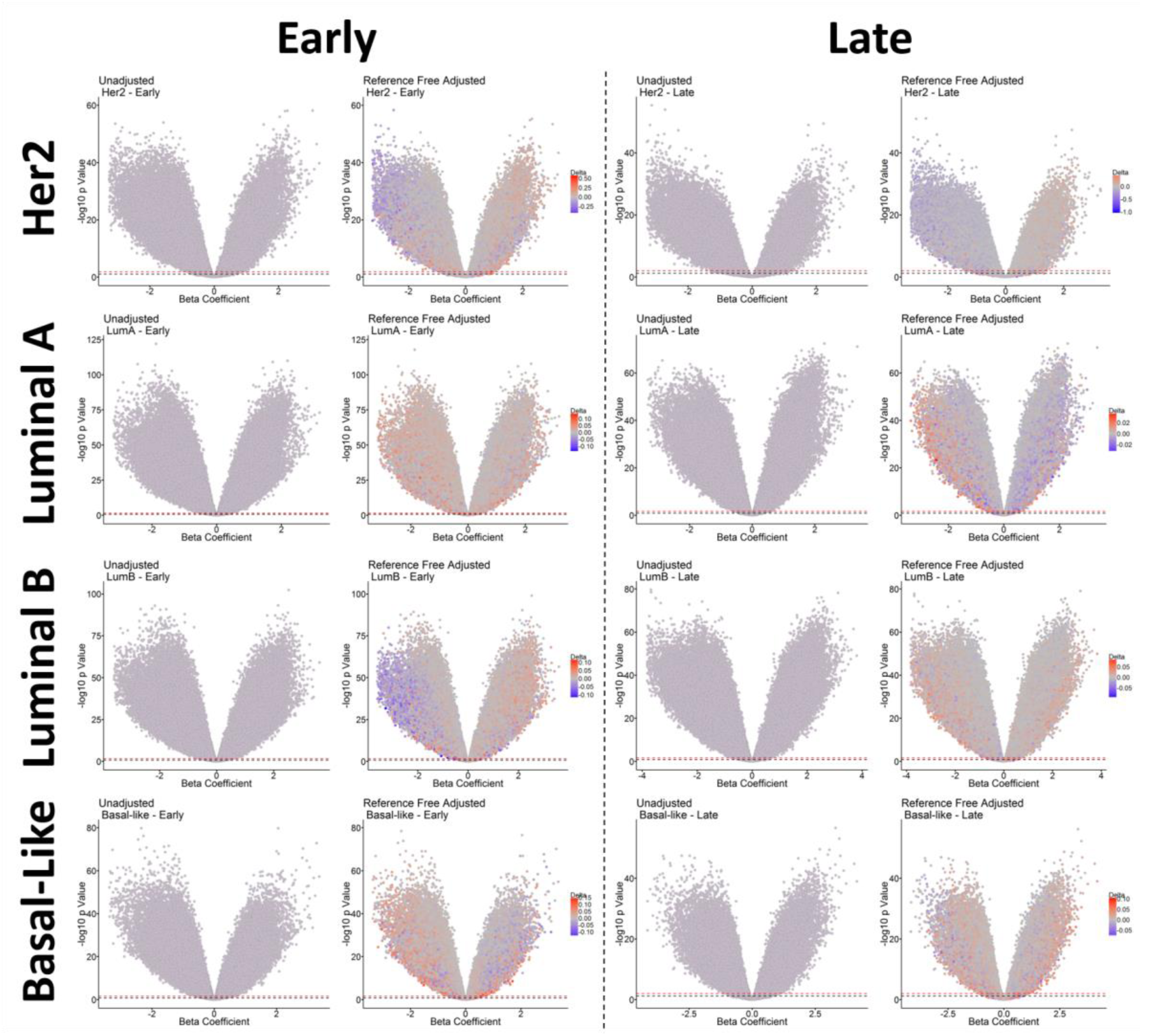
Volcano plots from the eight models. The left most panel in each model indicates unadjusted P values and the right panel indicates RefFree adjusted P values. Each point represents a CpG considered in the model and the color of the points represents the change in the beta coefficient following adjustment (delta value). The red lines indicate a Q value cutoff of 0.01 and the black lines indicate a Q value cutoff of 0.05.

**Supplementary Figure S3.**
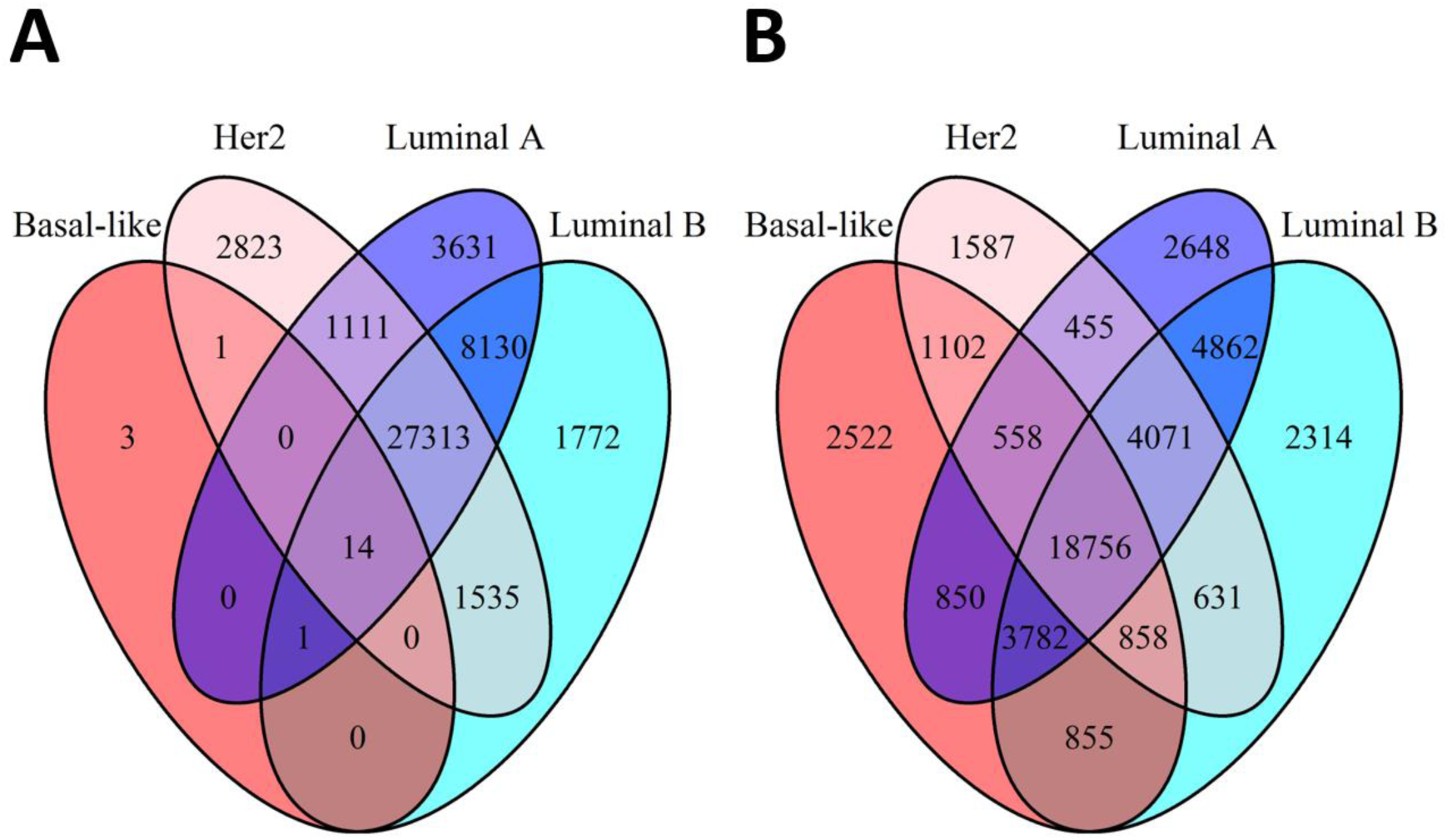
Venn diagram depicting overlapping Illumina annotation file UCSC regions between (A) early and (B) late stage tumors stratified by subtype. The regions consist of mappings relative to CpG island definitions (e.g. <Gene Name> N_Shore).

**Supplementary Figure S4.**
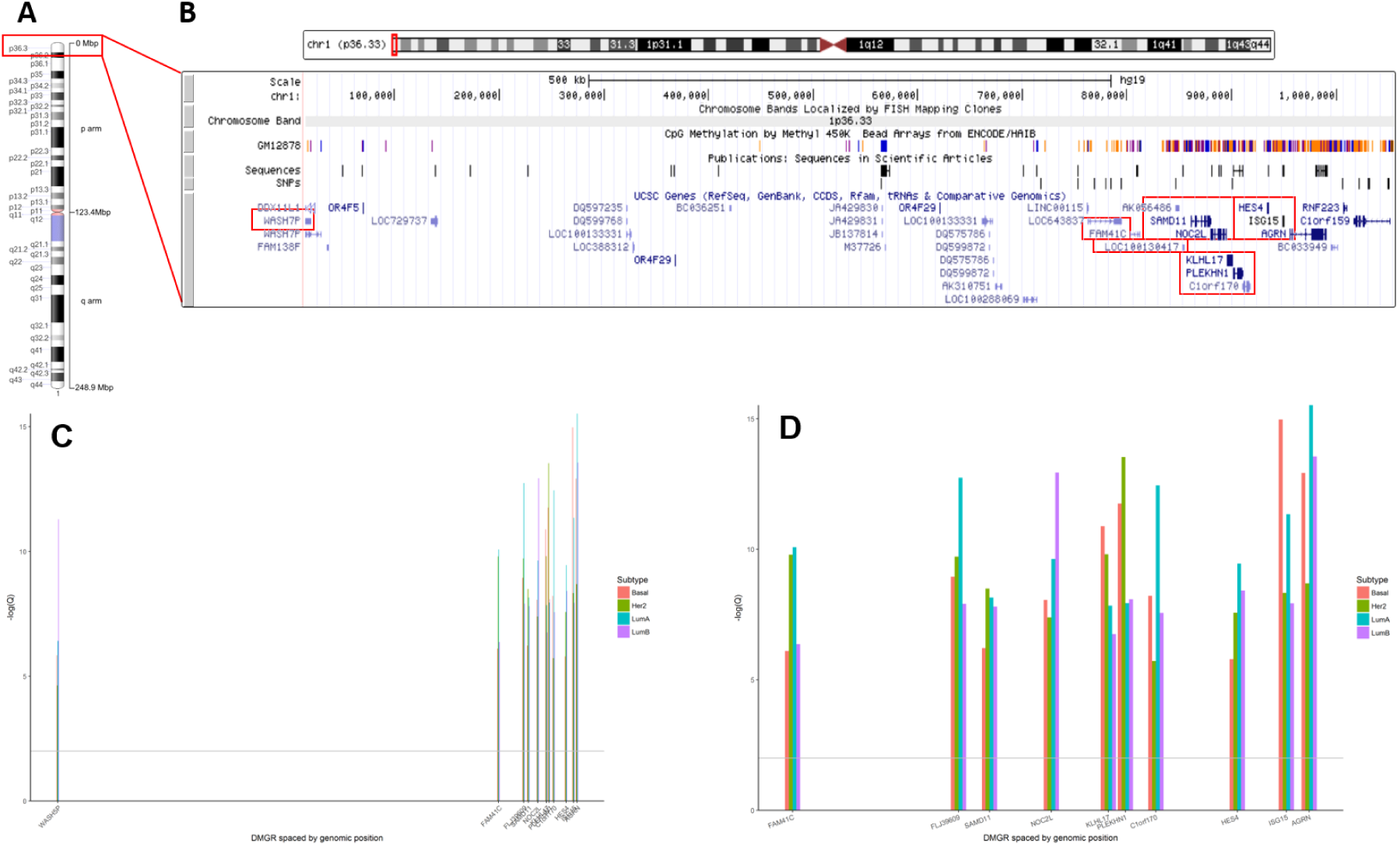
Diagram of chromosome 1. (A) The entire chromosome 1 with regions annotated. (B) A zoomed in view of chromosome 1p36.3 with each identified gene annotated on a track and highlighted in red boxes indicating a gene cluster between base pairs 868,071 - 1,056,116. (C) The negative log of the median Q-value for all CpG sites within each DMGR, stratified by PAM50 subtype and arranged along the x-axis according to genomic position reflected in panel B. (D) The negative log of the median Q-value for all CpG sites within each DMGR in the ten gene cluster (without WASH5P), stratified by PAM50 subtype and arranged along the x-axis according to genomic position reflected in panel B

**Supplementary Figure S5.**
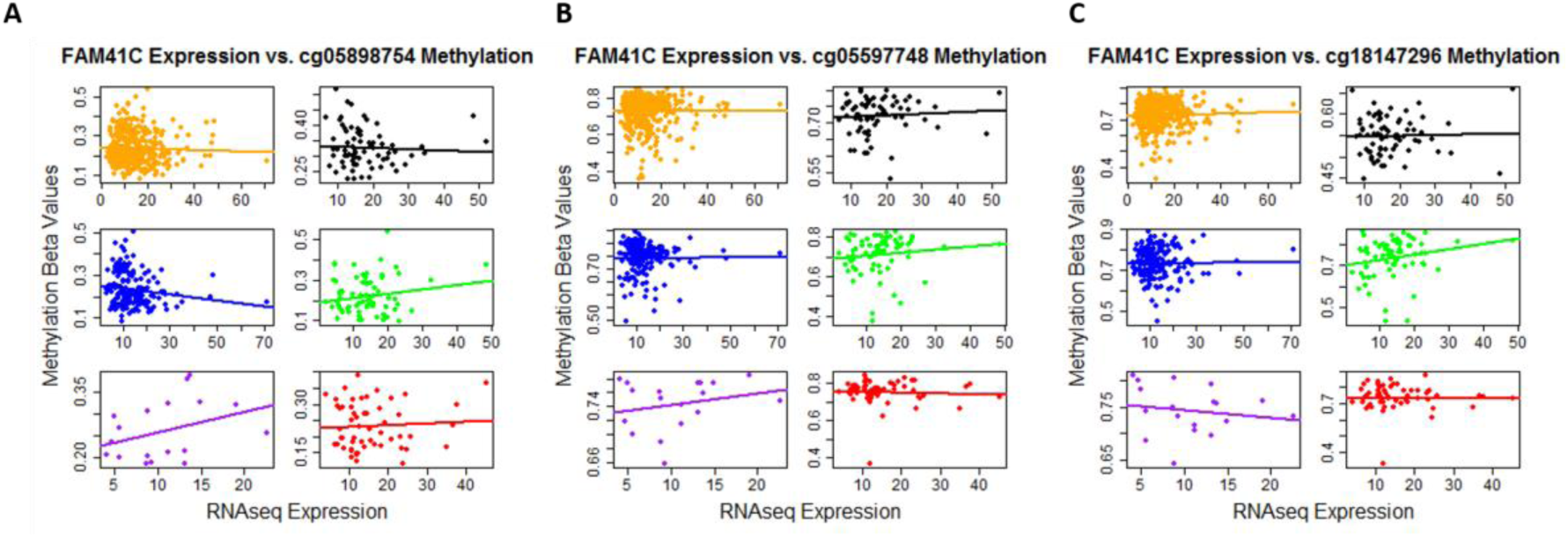
The relationship between differentially methylated CpG sites and FAM41C gene expression in early stage tumors and normal tissue with matched RNAseq samples stratified by PAM50 subtype. All tumors (orange), all normal tissue (black), Luminal A (blue), Luminal B (green), Her2 (purple), and Basal-like (red) are given in the different facets of the figure.

**Supplementary Figure S6.**
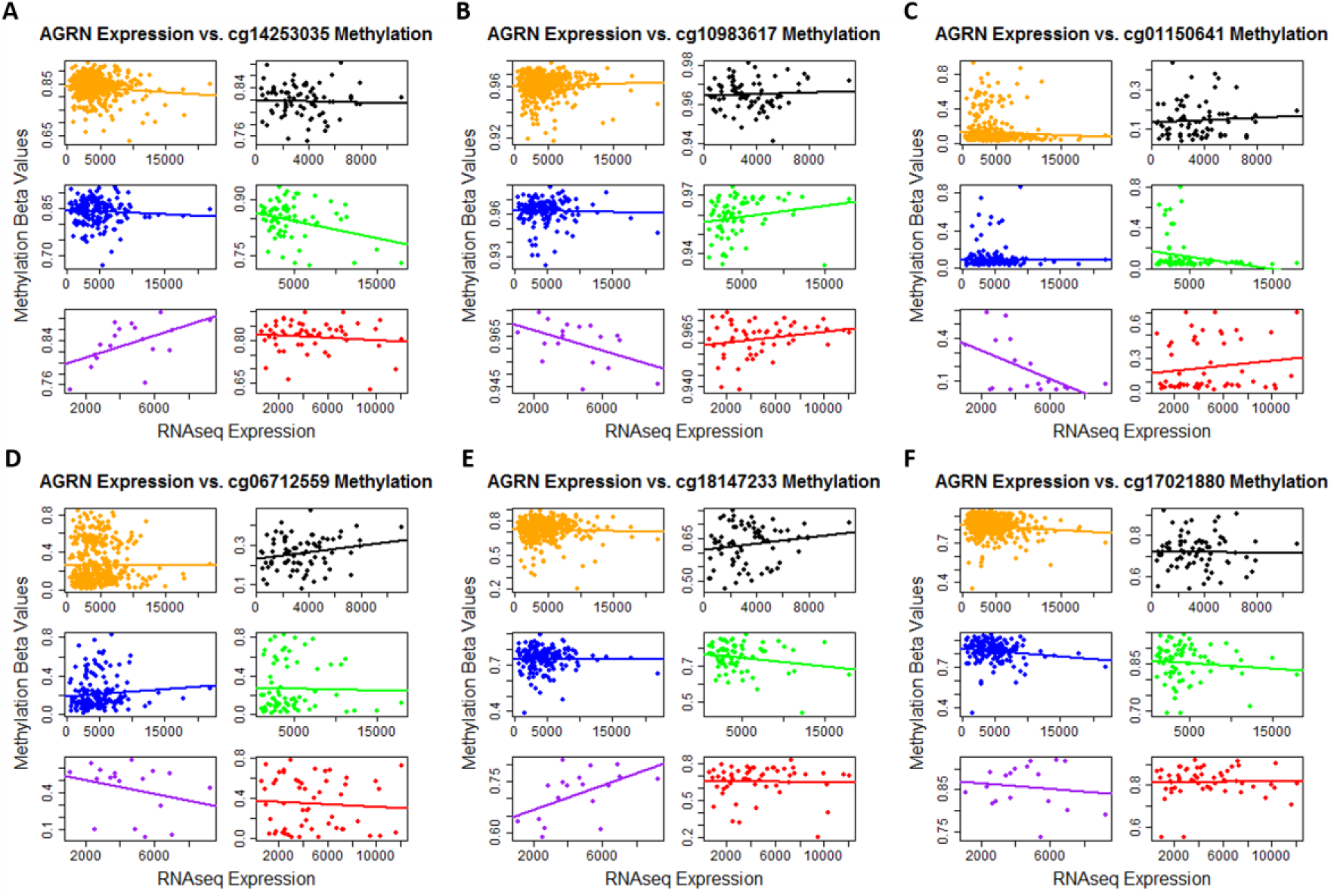
The relationship between differentially methylated CpG sites and AGRN gene expression in early stage tumors and normal tissue with matched RNAseq samples stratified by PAM50 subtype. All tumors (orange), all normal tissue (black), Luminal A (blue), Luminal B (green), Her2 (purple), and Basal-like (red) are given in the different facets of the figure.

**Supplementary Figure S7.**
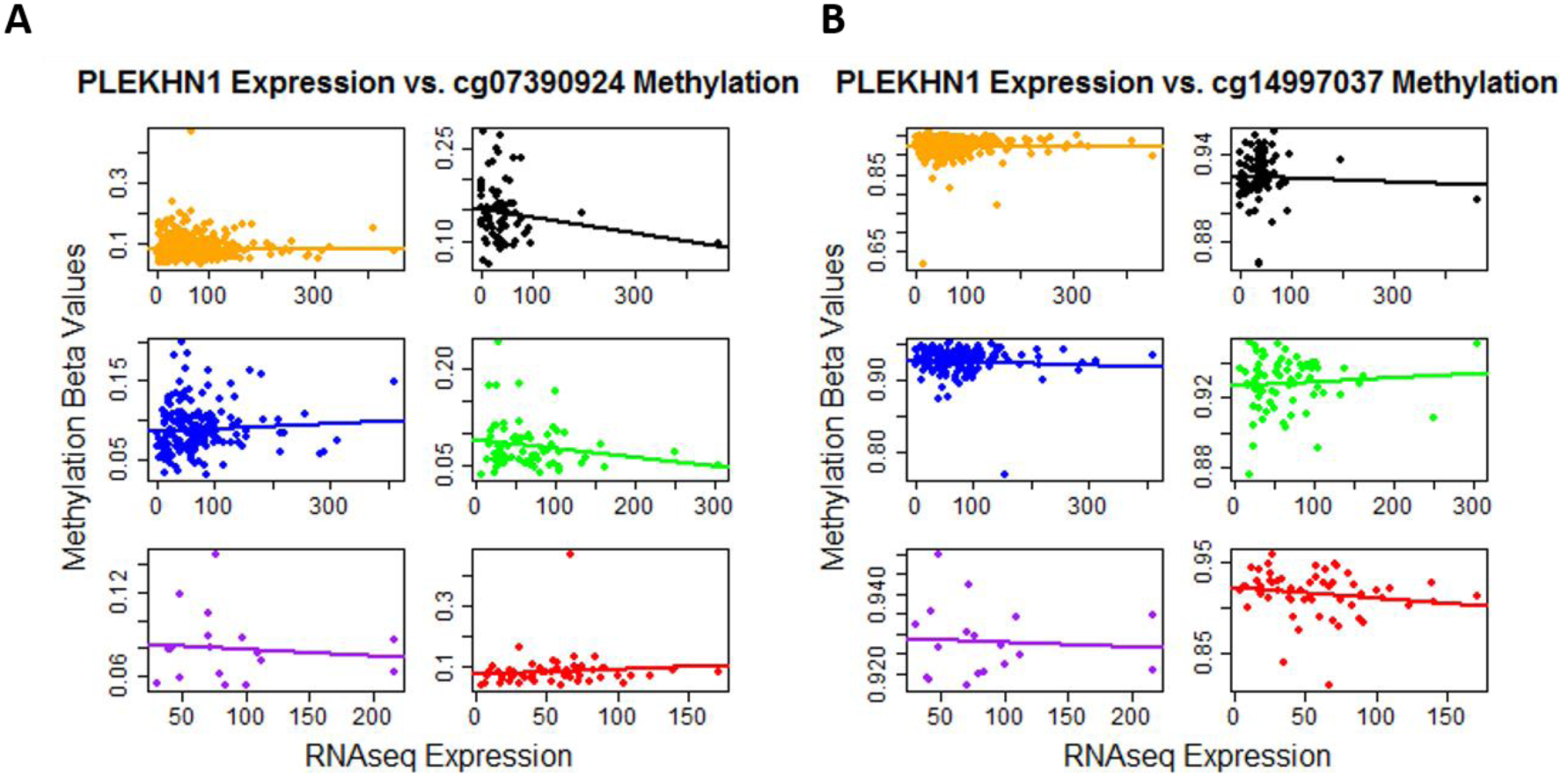
The relationship between differentially methylated CpG sites and PLEKHN1 gene expression in early stage tumors and normal tissue with matched RNAseq samples stratified by PAM50 subtype. All tumors (orange), all normal tissue (black), Luminal A (blue), Luminal B (green), Her2 (purple), and Basal-like (red) are given in the different facets of the figure.

**Supplementary Figure S8.**
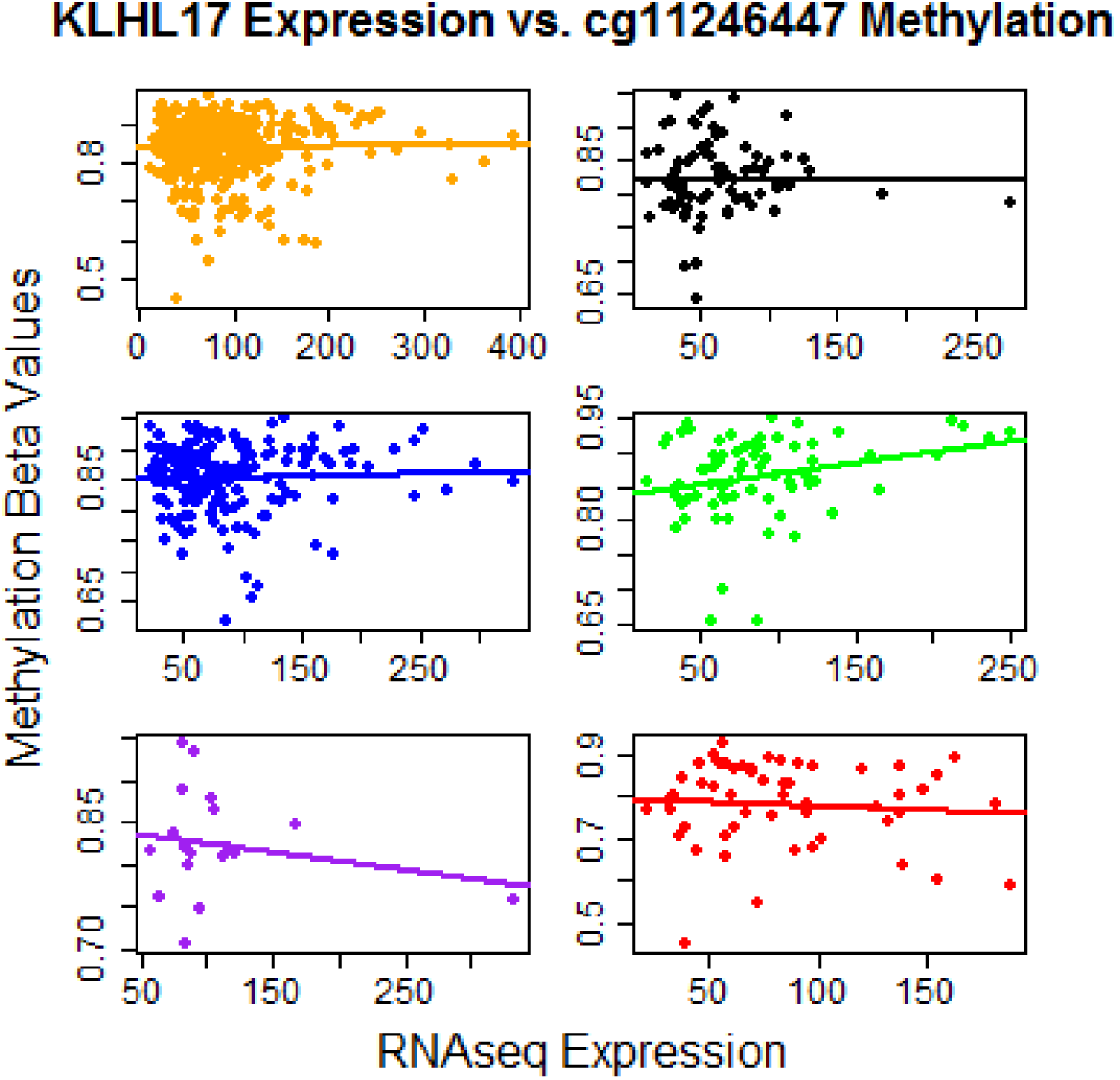
The relationship between differentially methylated CpG sites and KLHL17 gene expression in early stage tumors and normal tissue with matched RNAseq samples stratified by PAM50 subtype. All tumors (orange), all normal tissue (black), Luminal A (blue), Luminal B (green), Her2 (purple), and Basal-like (red) are given in the different facets of the figure.

**Supplementary Figure S9.**
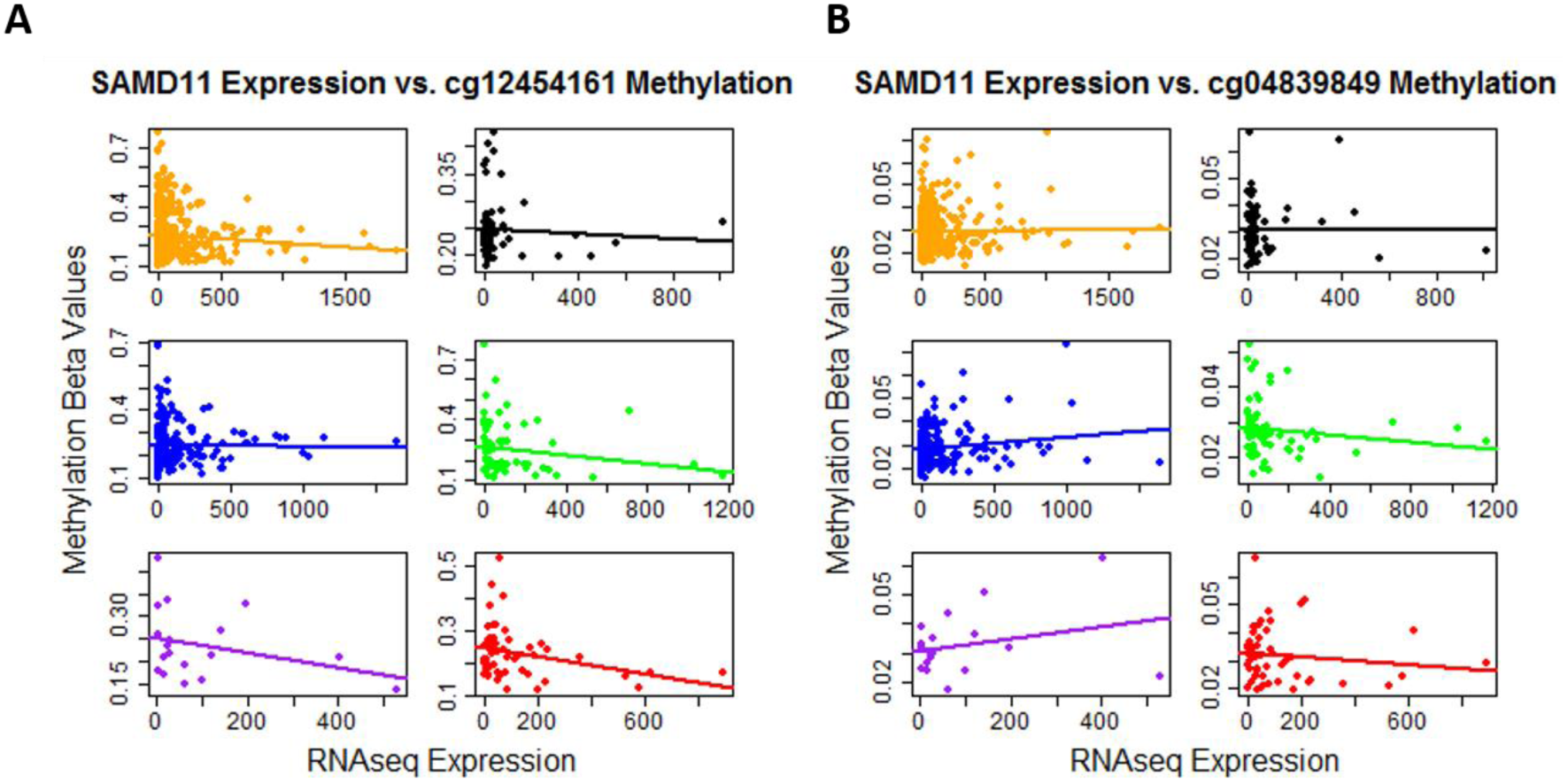
The relationship between differentially methylated CpG sites and SAMD11 gene expression in early stage tumors and normal tissue with matched RNAseq samples stratified by PAM50 subtype. All tumors (orange), all normal tissue (black), Luminal A (blue), Luminal B (green), Her2 (purple), and Basal-like (red) are given in the different facets of the figure.

**Supplementary Figure S10.**
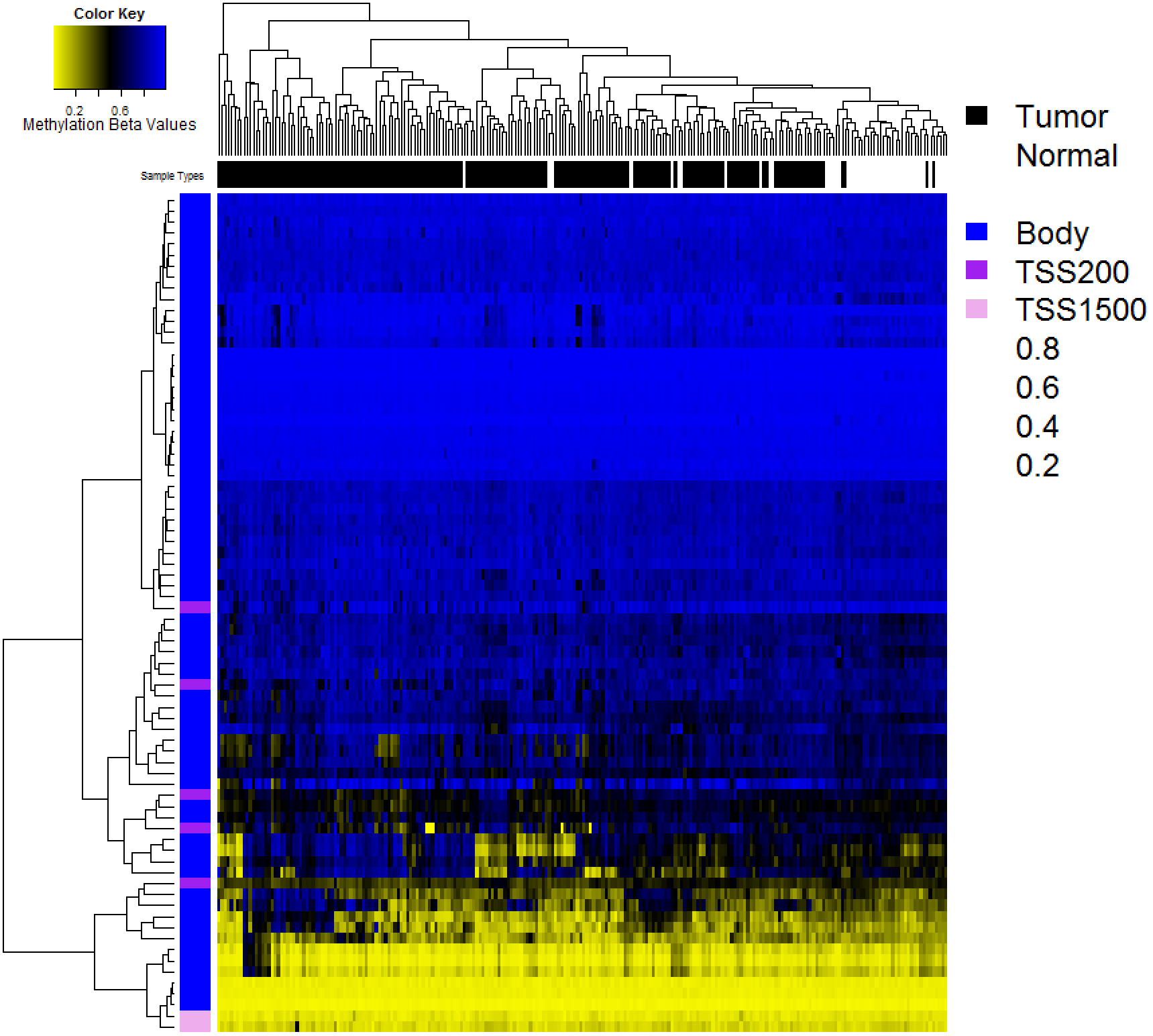
Results from the validation set (Fleischer et al 2014; GSE60185). Validation set raw (unadjusted) beta value heatmap of the significantly differentially methylated CpG sites in the common early stage differentially methylated gene regions (DMGRs) identified in the initial analysis. The genomic context is given in the vertical color bar (blue = gene body, dark pink = TSS200, light pink = TSS1500) and tumor vs. normal status is given in the horizontal color bar (black = tumor, white = normal tissue). In the heatmap, yellow indicates low methylation and blue indicates high methylation.

## SUPPLAMENTAL TABLES

Due to size limitations of this document and the size of the supplemental tables available for this manuscript, supplemental tables may be found at the following DOI link: DOI: 10.5281/zenodo.400247

## REFERENCES

1. Perou, CM, Sørlie T, Eisen, MB, van de Rijn, M, Jeffrey, SS, Rees, CA, Pollack, JR, Ross, DT, Johnsen, H, Akslen, LA, Fluge, O, Pergamenschikov, A, Williams, C, Zhu, SX, Lønning PE, Børresen-Dale, AL, Brown, PO, Botstein, D. Molecular portraits of human breast tumours. Nature. 2000 Aug 17;406(6797):747–752. PMID: 10963602

2. Vogelstein, B, Papadopoulos, N, Velculescu, VE, Zhou, S, Diaz, LA, Kinzler, KW. Cancer genome landscapes. Science. 2013 Mar 29;339(6127):1546–1558. PMCID: PMC3749880

3. Jones, PA, Baylin, SB. The fundamental role of epigenetic events in cancer. Nat Rev Genet. 2002 Jun;3(6):415–428. PMID: 12042769

4. Yang, X, Yan, L, Davidson, NE. DNA methylation in breast cancer. Endocr Relat Cancer. 2001 Jun;8(2):115–127. PMID: 11446343

5. Baylin, SB, Esteller, M, Rountree, MR, Bachman, KE, Schuebel, K, Herman, JG. Aberrant patterns of DNA methylation, chromatin formation and gene expression in cancer. Hum Mol Genet. 2001 Apr;10(7):687–692. PMID: 11257100

6. Fang, F, Turcan, S, Rimner, A, Kaufman, A, Giri, D, Morris, LGT, Shen, R, Seshan, V, Mo, Q, Heguy, A, Baylin, SB, Ahuja, N, Viale, A, Massague, J, Norton, L, Vahdat, LT, Moynahan, ME, Chan, TA. Breast cancer methylomes establish an epigenomic foundation for metastasis. Sci Transl Med. 2011 Mar 23;3(75):75ra25. PMCID: PMC3146366

7. Kamalakaran, S, Varadan, V, Giercksky Russnes, HE, Levy, D, Kendall, J, Janevski, A, Riggs, M, Banerjee, N, Synnestvedt, M, Schlichting, E, Karesen, R, Shama Prasada, K, Rotti, H, Rao, R, Rao, L, Eric Tang, M-H, Satyamoorthy, K, Lucito, R, Wigler, M, Dimitrova, N, Naume, B, Borresen-Dale, A-L, Hicks, JB. DNA methylation patterns in luminal breast cancers differ from non-luminal subtypes and can identify relapse risk independent of other clinical variables. Mol Oncol. 2011 Feb;5(1):77–92. PMID: 21169070

8. Sørlie T, Perou, CM, Tibshirani, R, Aas, T, Geisler, S, Johnsen, H, Hastie, T, Eisen, MB, van de Rijn, M, Jeffrey, SS, Thorsen, T, Quist, H, Matese, JC, Brown, PO, Botstein, D, Lønning PE, Børresen-Dale A-L. Gene expression patterns of breast carcinomas distinguish tumor subclasses with clinical implications. Proc Natl Acad Sci. 2001 Sep 11;98(19):10869–10874. PMID: 11553815

9. Cancer Genome Atlas Network. Comprehensive molecular portraits of human breast tumours. Nature. 2012 Oct 4;490(7418):61–70. PMCID: PMC3465532

10. Beca, F, Polyak, K. Intratumor Heterogeneity in Breast Cancer. Adv Exp Med Biol. 2016;882:169–189. PMID: 26987535

11. Yoshihara, K, Shahmoradgoli, M, Martinez, E, Vegesna, R, Kim, H, Torres-Garcia, W, Trevino, V, Shen, H, Laird, PW, Levine, DA, Carter, SL, Getz, G, Stemke-Hale, K, Mills, GB, Verhaak, RGW. Inferring tumour purity and stromal and immune cell admixture from expression data. Nat Commun. 2013;4:2612. PMCID: PMC3826632

12. Bloushtain-Qimron, N, Yao, J, Snyder, EL, Shipitsin, M, Campbell, LL, Mani, SA, Hu, M, Chen, H, Ustyansky, V, Antosiewicz, JE, Argani, P, Halushka, MK, Thomson, JA, Pharoah, P, Porgador, A, Sukumar, S, Parsons, R, Richardson, AL, Stampfer, MR, Gelman, RS, Nikolskaya, T, Nikolsky, Y, Polyak, K. Cell type-specific DNA methylation patterns in the human breast. Proc Natl Acad Sci U S A. 2008 Sep 16;105(37):14076–14081. PMCID: PMC2532972

13. Christensen, BC, Houseman, EA, Marsit, CJ, Zheng, S, Wrensch, MR, Wiemels, JL, Nelson, HH, Karagas, MR, Padbury, JF, Bueno, R, Sugarbaker, DJ, Yeh, R-F, Wiencke, JK, Kelsey, KT. Aging and environmental exposures alter tissue-specific DNA methylation dependent upon CpG island context. PLoS Genet. 2009 Aug;5(8):e1000602. PMCID: PMC2718614

14. Santagata, S, Thakkar, A, Ergonul, A, Wang, B, Woo, T, Hu, R, Harrell, JC, McNamara, G, Schwede, M, Culhane, AC, Kindelberger, D, Rodig, S, Richardson, A, Schnitt, SJ, Tamimi, RM, Ince, TA. Taxonomy of breast cancer based on normal cell phenotype predicts outcome. J Clin Invest. 2014 Feb;124(2):859–870. PMCID: PMC3904619

15. Koestler, DC, Christensen, B, Karagas, MR, Marsit, CJ, Langevin, SM, Kelsey, KT, Wiencke, JK, Houseman, EA. Blood-based profiles of DNA methylation predict the underlying distribution of cell types: a validation analysis. Epigenetics Off J DNA Methylation Soc. 2013 Aug;8(8):816–826. PMCID: PMC3883785

16. Houseman, EA, Kile, ML, Christiani, DC, Ince, TA, Kelsey, KT, Marsit, CJ. Reference-free deconvolution of DNA methylation data and mediation by cell composition effects. BMC Bioinformatics. 2016;17(1):259. PMID: 27358049

17. Houseman, EA, Kelsey, KT, Wiencke, JK, Marsit, CJ. Cell-composition effects in the analysis of DNA methylation array data: a mathematical perspective. BMC Bioinformatics. 2015 Mar 21;16:95. PMCID: PMC4392865

18. Houseman, EA, Ince, TA. Normal cell-type epigenetics and breast cancer classification: a case study of cell mixture-adjusted analysis of DNA methylation data from tumors. Cancer Inform. 2014;13(Suppl 4):53–64. PMCID: PMC4264613

19. Bagchi, A, Mills, AA. The Quest for the 1p36 Tumor Suppressor. Cancer Res. 2008 Apr 15;68(8):2551–2556. PMID: 18413720

20. Fleischer, T, Frigessi, A, Johnson, KC, Edvardsen, H, Touleimat, N, Klajic, J, Riis, ML, Haakensen, VD, Wärnberg F, Naume, B, Helland, A, Børresen-Dale, A-L, Tost, J, Christensen, BC, Kristensen, VN. Genome-wide DNA methylation profiles in progression to in situ and invasive carcinoma of the breast with impact on gene transcription and prognosis. Genome Biol. 2014;15(8):435. PMCID: PMC4165906

21. Titus, AJ, Way, GP, Johnson, KC, Christensen, BC. Analytical code for “Reference-free deconvolution of DNA methylation signatures identifies common differentially methylated gene regions on 1p36 across breast cancer subtypes.” 2017 Mar 10; Available from: https://zenodo.org/badge/latestdoi/45754471

22. Fang, F, Turcan, S, Rimner, A, Kaufman, A, Giri, D, Morris, LGT, Shen, R, Seshan, V, Mo, Q, Heguy, A, Baylin, SB, Ahuja, N, Viale, A, Massague, J, Norton, L, Vahdat, LT, Moynahan, ME, Chan, TA. Breast Cancer Methylomes Establish an Epigenomic Foundation for Metastasis. Sci Transl Med. 2011 Mar 23;3(75):75ra25. PMID: 21430268

23. Orsetti, B, Nugoli, M, Cervera, N, Lasorsa, L, Chuchana, P, Rouge, C, Ursule, L, Nguyen, C, Bibeau, F, Rodriguez, C, Theillet, C. Genetic profiling of chromosome 1 in breast cancer: mapping of regions of gains and losses and identification of candidate genes on 1q. Br J Cancer. 2006 Nov 20;95(10):1439–1447. PMCID: PMC2360604

24. Munn, KE, Walker, RA, Varley, JM. Frequent alterations of chromosome 1 in ductal carcinoma in situ of the breast. Oncogene. 1995 Apr 20;10(8):1653– 1657. PMID: 7731721

25. Mamessier, E, Song, JY, Eberle, FC, Pack, S, Drevet, C, Chetaille, B, Abdullaev, Z, Adelaide, J, Birnbaum, D, Chaffanet, M, Pittaluga, S, Roulland, S, Chott, A, Jaffe, ES, Nadel, B. Early lesions of follicular lymphoma: a genetic perspective. Haematologica. 2014 Mar;99(3):481–488. PMCID: PMC3943311

26. Bieche, I, Champeme, MH, Lidereau, R. Loss and gain of distinct regions of chromosome 1q in primary breast cancer. Clin Cancer Res Off J Am Assoc Cancer Res. 1995 Jan;1(1):123–127. PMID: 9815894

27. Curtis, C, Shah, SP, Chin S-F, Turashvili, G, Rueda, OM, Dunning, MJ, Speed, D, Lynch, AG, Samarajiwa, S, Yuan, Y, Graf, S, Ha, G, Haffari, G, Bashashati, A, Russell, R, McKinney, S, Langerod, A, Green, A, Provenzano, E, Wishart, G, Pinder, S, Watson, P, Markowetz, F, Murphy, L, Ellis, I, Purushotham, A, Borresen-Dale A-L, Brenton, JD, Tavare, S, Caldas, C, Aparicio, S. The genomic and transcriptomic architecture of 2,000 breast tumours reveals novel subgroups. Nature. 2012 Apr 18;486(7403):346–352. PMCID: PMC3440846

28. White, PS, Thompson, PM, Gotoh, T, Okawa, ER, Igarashi, J, Kok, M, Winter, C, Gregory, SG, Hogarty, MD, Maris, JM, Brodeur, GM. Definition and characterization of a region of 1p36.3 consistently deleted in neuroblastoma. Oncogene. 2005 Apr 14;24(16):2684–2694. PMID: 15829979

29. Attiyeh, EF, London, WB, Mossé YP, Wang, Q, Winter, C, Khazi, D, McGrady, PW, Seeger, RC, Look, AT, Shimada, H, Brodeur, GM, Cohn, SL, Matthay, KK, Maris, JM. Chromosome 1p and 11q Deletions and Outcome in Neuroblastoma. N Engl J Med. 2005 Nov 24;353(21):2243–2253. PMID: 16306521

30. Caren, H, Ejeskar, K, Fransson, S, Hesson, L, Latif, F, Sjoberg R-M, Krona, C, Martinsson, T. A cluster of genes located in 1p36 are down-regulated in neuroblastomas with poor prognosis, but not due to CpG island methylation. Mol Cancer. 2005 Mar 1;4(1):10. PMCID: PMC554762

31. Carén H, Fransson, S, Ejeskär K, Kogner, P, Martinsson, T. Genetic and epigenetic changes in the common 1p36 deletion in neuroblastoma tumours. Br J Cancer. 2007 Nov 19;97(10):1416–1424. PMCID: PMC2360241

32. Henrich K-O, Schwab, M, Westermann, F. 1p36 tumor suppression--a matter of dosage? Cancer Res. 2012 Dec 1;72(23):6079–6088. PMID: 23172308

33. Lefebvre, C, Bachelot, T, Filleron, T, Pedrero, M, Campone, M, Soria J-C, Massard, C, Levy, C, Arnedos, M, Lacroix-Triki, M, Garrabey, J, Boursin, Y, Deloger, M, Fu, Y, Commo, F, Scott, V, Lacroix, L, Dieci, MV, Kamal, M, Dieras, V, Goncalves, A, Ferrerro J-M, Romieu, G, Vanlemmens, L, Mouret Reynier, M-A, Thery J-C, Le Du, F, Guiu, S, Dalenc, F, Clapisson, G, Bonnefoi, H, Jimenez, M, Le Tourneau, C, Andre, F. Mutational Profile of Metastatic Breast Cancers: A Retrospective Analysis. PLoS Med. 2016 Dec;13(12):e1002201. PMCID: PMC5189935

34. Stoeck, A, Lejnine, S, Truong, A, Pan, L, Wang, H, Zang, C, Yuan, J, Ware, C, MacLean, J, Garrett-Engele, PW, Kluk, M, Laskey, J, Haines, BB, Moskaluk, C, Zawel, L, Fawell, S, Gilliland, G, Zhang, T, Kremer, BE, Knoechel, B, Bernstein, BE, Pear, WS, Liu, XS, Aster, JC, Sathyanarayanan, S. Discovery of biomarkers predictive of GSI response in triple-negative breast cancer and adenoid cystic carcinoma. Cancer Discov. 2014 Oct;4(10):1154–1167. PMCID: PMC4184927

35. Burks, J, Reed, RE, Desai, SD. Free ISG15 triggers an antitumor immune response against breast cancer: a new perspective. Oncotarget. 2015 Mar 30;6(9):7221–7231. PMCID: PMC4466680

36. Andersen, JB, Hassel, BA. The interferon regulated ubiquitin-like protein, ISG15, in tumorigenesis: friend or foe? Cytokine Growth Factor Rev. 2006 Dec;17(6):411–421. PMID: 17097911

37. Xu, L, Tan, AC, Winslow, RL, Geman, D. Merging microarray data from separate breast cancer studies provides a robust prognostic test. BMC Bioinformatics. 2008 Feb 27;9:125. PMCID: PMC2409450

38. Rodriguez-Martinez, A, Alarmo E-L, Saarinen, L, Ketolainen, J, Nousiainen, K, Hautaniemi, S, Kallioniemi, A. Analysis of BMP4 and BMP7 signaling in breast cancer cells unveils time-dependent transcription patterns and highlights a common synexpression group of genes. BMC Med Genomics. 2011 Nov 25;4:80. PMCID: PMC3229454

39. Hansen, KD, Fortin, JP. Minfi tutorial. BioC2014. 2014;

40. Chen, Y, Lemire, M, Choufani, S, Butcher, DT, Grafodatskaya, D, Zanke, BW, Gallinger, S, Hudson, TJ, Weksberg, R. Discovery of cross-reactive probes and polymorphic CpGs in the Illumina Infinium HumanMethylation450 microarray. Epigenetics Off J DNA Methylation Soc. 2013 Feb;8(2):203–209. PMCID: PMC3592906

41. Wilhelm-Benartzi, CS, Koestler, DC, Karagas, MR, Flanagan, JM, Christensen, BC, Kelsey, KT, Marsit, CJ, Houseman, EA, Brown, R. Review of processing and analysis methods for DNA methylation array data. Br J Cancer. 2013 Sep 17;109(6):1394–1402. PMCID: PMC3777004

42. Edge, S, Byrd, D, Compton, C, Fritz, A, Greene, F, Trotti, A, editors. AJCC cancer staging manual. 7th ed. New York, NY: Springer; 2010.

43. Brunet J-P, Tamayo, P, Golub, TR, Mesirov, JP. Metagenes and molecular pattern discovery using matrix factorization. Proc Natl Acad Sci U S A. 2004 Mar 23;101(12):4164–4169. PMCID: PMC384712

44. Dabney, A, Storey, J. qvalue: Q-value estimation for false discovery rate control. R Package Version 1430.

45. Huang, DW, Sherman, BT, Lempicki, RA. Bioinformatics enrichment tools: paths toward the comprehensive functional analysis of large gene lists. Nucleic Acids Res. 2009 Jan;37(1):1–13. PMCID: PMC2615629

46. Huang, DW, Sherman, BT, Lempicki, RA. Systematic and integrative analysis of large gene lists using DAVID bioinformatics resources. Nat Protoc. 2009;4(1):44–57. PMID: 19131956

47. Ashburner, M, Ball, CA, Blake, JA, Botstein, D, Butler, H, Cherry, JM, Davis, AP, Dolinski, K, Dwight, SS, Eppig, JT, Harris, MA, Hill, DP, Issel-Tarver, L, Kasarskis, A, Lewis, S, Matese, JC, Richardson, JE, Ringwald, M, Rubin, GM, Sherlock, G. Gene ontology: tool for the unification of biology. The Gene Ontology Consortium. Nat Genet. 2000 May;25(1):25–29. PMCID: PMC3037419

48. Kanehisa, M, Goto, S. KEGG: kyoto encyclopedia of genes and genomes. Nucleic Acids Res. 2000 Jan 1;28(1):27–30. PMCID: PMC102409

49. Huang, DW, Sherman, BT, Lempicki, RA. Systematic and integrative analysis of large gene lists using DAVID bioinformatics resources. Nat Protoc. 2009;4(1):44–57. PMID: 19131956

50. Gao, J, Aksoy, BA, Dogrusoz, U, Dresdner, G, Gross, B, Sumer, SO, Sun, Y, Jacobsen, A, Sinha, R, Larsson, E, Cerami, E, Sander, C, Schultz, N. Integrative analysis of complex cancer genomics and clinical profiles using the cBioPortal. Sci Signal. 2013 Apr 2;6(269):pl1. PMCID: PMC4160307

